# Properties of the full random effect modelling approach with missing covariates

**DOI:** 10.1101/656470

**Authors:** Joakim Nyberg, E. Niclas Jonsson, Mats O. Karlsson, Jonas Häggström, members of the HBGDki Community

**Affiliations:** Pharmetheus AB, Uppsala, Sweden; Department of Pharmaceutical Biosciences, Uppsala University, Uppsala, Sweden; Mtek Sciences, Stockholm, Sweden

**Keywords:** FREM, full modelling approach, height-for-age Z-score, global health

## Abstract

Two full model approaches was compared with respect to their ability to handle missing covariate information. The reference data analysis approach was the full model method in which the covariate effects are estimated conventionally using fixed effects, and missing covariate data is imputed with the median of the non-missing covariate information. This approach was compared to a novel full model method which treats the covariate data as observed data and estimates the covariates as random effects. A consequence of this way of handling the covariates is that no covariate imputation is required and that any missingness in the covariates is handled implicitly. The comparison between the two analysis methods was based on simulated data from a model of height for age z-scores as a function of age. Data was simulated with increasing degrees of randomly missing covariate information (0-90%) and analyzed using each of the two analysis approaches. Not surprisingly, the precision in the parameter estimates from both methods decreased with increasing degrees of missing covariate information. However, while the bias in the parameter estimates increased in a similar fashion for the reference method, the full random effects approach provided unbiased estimates for all degrees of covariate missingness.

## 1 Introduction

Covariates are observable predictors that may be included in, for example, nonlinear mixed effects models (NLMEM), to reduce unexplained variability. The identification and estimation of the coefficients for covariates can be done in many different ways. Very broadly, such methods can be classified as either selection methods or full model approaches. Simply put, selection methods use the observed data to both select the covariates to include in the model as well as to estimate their coefficients. The decision to select a particular covariate for inclusion in the model is typically based on an objective criterion, such as a p-value. A typical example of a selection method is stepwise linear regression. There are well known weaknesses with selection methods, e.g. the risk for selection bias, that the order of inclusion of the covariates may bias the results and a sensitivity to correlations between the covariates. The advantages include parsimonious final models and relative simplicity. In contrast to selection methods, full model approaches include a pre specified set of covariates in the model regardless of statistical significance. The pre specification effectively avoids selection bias and makes inference more correct. However, full model approaches still has issues with correlations and, with non-linear models without analytical solutions, there may be numerical problems if the number of covariates is too large. An example in the field of pharmacometrics is the full model approach suggested by Gastonguay (1). This approach includes a number of steps from pre specification to inference, including a "data reduction step", which involves removing covariates from the pre specified set in case of too high correlations(3, 4).

A common challenge with all the above methods is how to handle missing covariates. There are many ways to handle missing covariates and depends on if the data are missing at random or not. If data are not missing at random, strategies such as building models for the mechanism of missing data are viable ways to handle the missing information. On the other hand, if covariate data are missing at random or completely at random, other techniques may be used, for example mean or median imputation, or more sophisticated methods such as multivariate multiple imputation (2).

In the present paper we will discuss a new full model method - the full random effect model (full random effects model (FREM)) approach (6–9). In this method the covariates are treated as observed data points and are modelled as random effects instead of being treated as error free explanatory variables whose impact of the model is estimated through fixed effect parameters, i.e. a full fixed effect model (full fixed effects model (FFEM)). Since covariances between random effects can be explicitly acknowledged in NLMEM, correlated covariates should be less of a problem with the FREM approach.

FREM implicitly handles missing data since non-observed covariates are simply not present in the vector of observed data in the dataset and therefore not used when estimating the FREM parameters. On the other hand, with FREM, the variance-covariance matrix of the parameters and covariates are estimated for the whole population. This means that missing values are still implicitly informed by the correlation to other covariates and dependent variables.

In this paper the FREM methodology will be investigated using a global health data set which was pooled from five separate studies. Because of its ability to handle missing covariate information, the FREM approach should be particularly useful for data from global health studies since such studies often contain a high number of highly correlated covariates and often have a large degree of missing covariate information. The latter is especially true if the data set is pooled from different studies, i.e. when it is likely to not be a complete overlap or comparability of covariates between the studies involved.

A FREM model was built using the public health data set and then used to further investigate the properties of FREM methodology to handle missing data compared to a simple mean imputation implemented in an FFEM. The performance of FREM versus FFEM are illustrated by visualizing bias and precision in a simulation and re-estimation study in which the degrees of missing covariate information is varied between 0% and 90%.

The paper is divided into a methods section where the FREM methodology is described in detail (Section 2) followed by a data section where the global health data is described together with the strategy for building the model using the FREM method (Section 3). This is followed by a section with the description of the simulation study used to investigate the missing covariate data properties of the FREM (Section 4). The results section first presents the FREM that was developed to describe the global health data (Section 5.1) followed by the results of the missing covariate properties of FREM (Section 5.2). The final section discusses the FREM results, implications and future perspectives (Section 6).

## 2 Full random effects model

A NLMEM for subject *i* and time point *j* without any covariates is described as

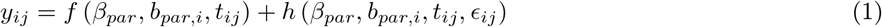

where *f* (.) is a non-linear function with respect to the structural parameters *β*_*par*_ and the random effects corresponding to the inter-individual variability (IIV), *b*_*par,i*_ ∼ 𝒩 (0, *D*_*par*_), *t*_*ij*_ is the independent variable (e.g. observation time), *h* (.) is a non-linear function describing how the individual observations differs from the model, i.e. the residual error function. *ϵ*_*ij*_ is the random effect corresponding to the residual unexplained variability (RUV), *ϵ*_*ij*_ ∼ 𝒩 (0, Σ). *D*_*par*_ and Σ are the covariance matrices of the individual random effects (*b*_*par,i*_) and the residual errors (*ϵ*_*ij*_) respectively.

A multi-response NLMEM is defined as a NLMEM with several types of observations. For example in drug development modelling this may be drug concentrations which are driving an outcome variable dependent on the drug concentration model. A multi-response NLMEM of *K* responses can be defined as

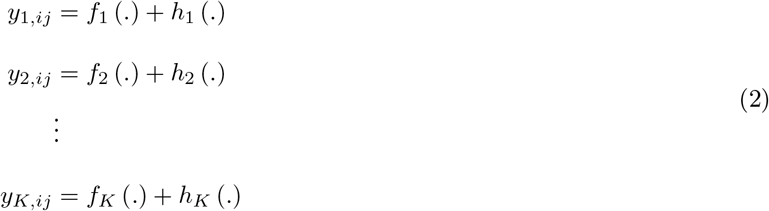

where each *k* response can have different parameters, models and independent variables.

The FREM definition make use of the multi-response nature of a NLMEM since all covariates are treated as separate response models without (possibly) any common parameters with the structural model but that are instead linked via the covariances of the individual random effects:

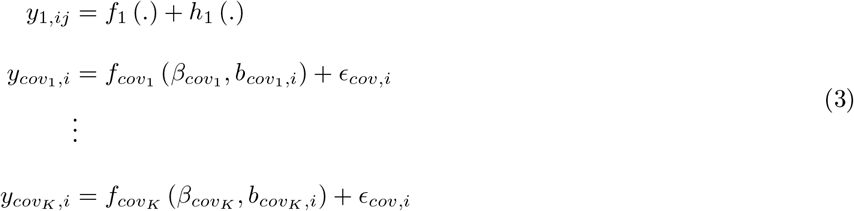

where 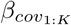 and 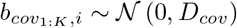 are the parameters describing the *K* covariates, *ϵ*_*cov,i*_ is a residual which variance (Σ_*cov*_) could be fixed to a value close to machine epsilon, (Σ_*cov*_ → *ϵ*) if the covariates are assumed measured without measurement error. The equation above describes one response (if not counting the covariates) but the methodology is not limited to only one response, the complete set of different types of observations (see Equation 2) can be linked within the FREM approach. Note that, so far, the *K* covariates are assumed time-invariant, hence the index *i* for the covariates. In the simplest form, the model for the *k* covariate is described as

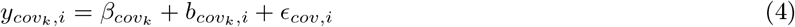

hence the 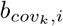 is estimated to be close to the difference between the observed covariate value and the mean of the covariate 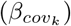.

In the case of complete data, this form, perhaps non-intuitively, does not assume any distributional assumption of the covariate distribution. In fact both continuous and categorical covariates can be parametrized in the same way. This is not surprising since the 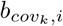 is the empirical Bayes estimate (EBE), estimated with a negligible residual error. For illustration, an example describing this is presented below.

### 2.1 Motivating example of a bi-modal covariate SEX

Assume that the sex of 100 subjects are observed as a covariate. 70 of the subjects are males and 30 subjects are females. If males are encoded as 1 and females as 2 the mean of the covariate (or the estimated unbiased mean) is 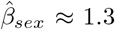. Furthermore, the random effect *b*_*sex,i*_ for each subject *i* is estimated to 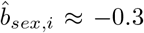 for males and 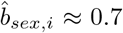 for females. Hence the underlying assumption of normality in the random effect *b*_*sex,i*_ is not distorting the bi-modal nature of the sex covariate in the estimated covariate distribution.

### 2.2 Linking FREM to a full fixed effect model

A feature of FREM is that it can be transformed to any combination of a fixed effect covariate model, i.e. a NLMEM with covariates handled as fixed effects. This implies that any FREM model with *K* covariates can be expressed as any combination of the *K*^2^ possible fixed effect covariate models without re-estimating the parameters. This is done by recalculating the parameter coefficients (*β*_*par*_) and the parameter variance (*D*_*par*_) by using the fact that *b*_*par*_ conditional on *b*_*cov*_ = *a*_*cov*_ (where *a*_*cov,i*_ is known) is a multivariate normal with mean

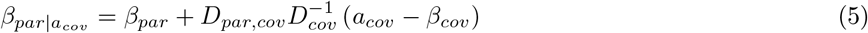

and variance

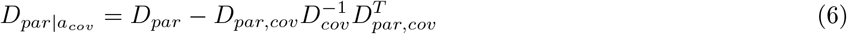

where the full inter individual covariance matrix *D* is defined as

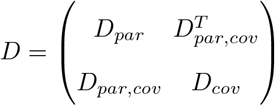

Note that it is only the rightmost part in Equation 5 that is dependent on the actual value of the covariate through *a*_*cov*_ which enables efficient calculations of the conditional parameters for different covariate values. Another remark is that the rows and columns of covariates that should be excluded in the fixed effect model are removed from *D*_*cov*_ and *D*_*par,cov*_ but their correlations to other included covariates will still influence the 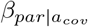 and 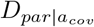.

### 2.3 Implicit handling of missing data

Since the FREM approach uses all the information about the correlation of covariates and parameters, random effects for missing covariates are implicitly estimated. As an illustration; assume two highly correlated covariates. If one of two covariate values from one subject is missing the correlation information about the non-missing covariate observation and the parameters will inform the estimation of the missing covariates even though it is not observed. Opposite to the complete data case, the distribution of the missing covariate is influenced by the parametrization of the covariate model. Compared to the example above (see Section 2.1) it is now possible that the correlations with the other covariates and the parameters gives a random effects estimate of e.g. 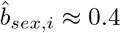. In this case the estimate indicates that it is more likely that the subject is a female than a male even though the estimated imputed value is neither female (2) nor male (1) but somewhere in between 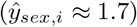.

Similar issues might arise when estimating missing continuous distributed covariates. For example a log-normally distributed covariate will be imputed based on normal assumptions if the simple additive form of the covariate model is used (see Equation 4). A way to avoid this is to either log-transform the data of the covariate or to change the covariate model *f*_*cov*_ (*β*_*cov*_, *b*_*cov,i*_) to a log-normal model. However, both of these modifications have impact on how the covariate-parameter model are interpreted as described in the next section.

### 2.4 Interpretation of fixed effects covariate-parameter relationships

The basic NLMEM equation(Equation 7) can be further expanded by defining the functional form of the relation between the inter-individual random effects (*b*_*par,i*_) and the structural parameters *β*_*par*_

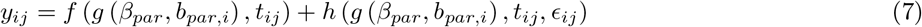

where *g* (.) is a (potentially) non-linear function. Examples of distributions of *g* (.) are

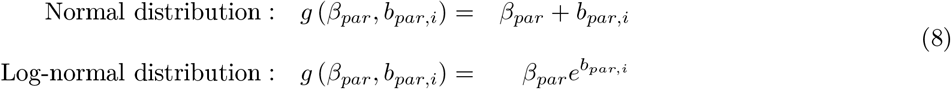

Since the relationship with the conditional normal distribution mean is linear with respect to the random effect (see Equation 5) the parameter-covariate relationship will be a function of the parameter distribution *g* (.) and the covariate model parametrization *f*_*cov*_ (.) (or covariate data transformation). For example a normal covariate model (*f*_*cov*_ = *β*_*cov*_ + *b*_*cov,i*_ + *ϵ*_*cov*_) with an exponential parameter model (*g*_*par*_ = *β*_*par*_*e*^*b*_*par,i*_^) yields a additive parameter-covariate relationship on the log scale:

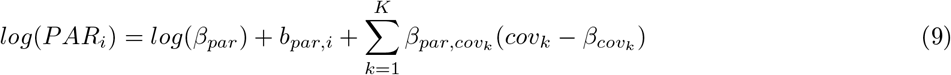

where the covariate contribution are calculated according to Equation 5.

Similarly, a parameter on the normal scale, will yield a parameter-covariate relationship on the normal scale:

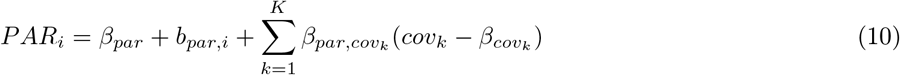

Most of the parameter-covariate relationships that are available in the full fixed effect modelling approach can be implemented (by covariate and/or parameter transformations) in FREM but some are only possible to approximate. For example, the log-normal proportional effect is not trivial to derive with the FREM approach, i.e. 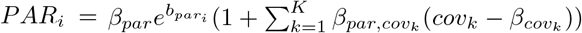.

### 2.5 Simulating with a FREM model

Data can be directly simulated using a FREM together with simulations of covariates since the covariate distributions are explicitly encoded in a FREM. However, when simulating covariates, there is no guarantee that the observed covariate distributions are kept. One way to ensure that e.g. sex is a dichotomous variable is either to round the samples into a dichotomous distribution or using transformations of normal samples into binomial samples via the covariate parametrization function *f*_*cov*_ (.). A feature when simulating data and covariates directly in FREM is that covariates are simulated using their estimated covariances, both to the data (via the parameter variances) as well as between covariates.

The alternative way to simulate data is to transform the model into a FFEM. After the transformation, it is trivial to simulate data using the observed covariates and the FFEM. Note though that in order to simulate using the observed covariates, a FFEM for each missing covariate pattern is needed since the covariate coefficients and IIV (see Equation 5) are dependent on which covariates that are missing. Hence, to correctly simulate using *K* covariates as many as *K*^2^ FFEMs might be needed, depending on if all combinations of missing covariates are present or not. On the other hand if several subjects have the same missing covariate pattern they can all be simulated using the same realized FFEM, which might decrease the computational burden when simulating large studies.

The later technique is useful when e.g. producing visual predictive checks (VPCs) (ref). Then one often use the realized design and observed covariates and it is important to acknowledge each subjects missing pattern.

### 2.6 Relate covariate effects to explained variability

Since the FREM methodology allows for any combination of the included covariates, a quantity that is of interest is the explained variability of a single covariate effect. This is a concept that describes how much a particular covariate (or a combination of covariates) explains of the total variability in the observed data (or in any transformation of the parameters, i.e. secondary parameters). The variability in the observed data can be divided into the variability due to IIV, RUV and covariates and it is trivial within the FREM methodology to scale the *D*_*par*_ by either 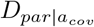 or using the EBEs to calculate the variability in the secondary parameters. Similar calculations can be applied using all covariate effects but without IIV and hence the total explainable variability by the observed covariates can be assessed. Other standard plots as e.g. forest plot (ref) are also possible to generate using a FREM.

## 3 COHORTs study

The COHORTs study (10) consist of five different studies in Brazil, Guatemala, India, Philippines and South Africa where children were followed for up to 19 years and longitudinal height for age Z-score (HAZ) observations were collected among other variables. The sampling design differed between the sites both in number of observations, duration of observations as well as observations times as is illustrated in Figure 1 and Table 1. The collected information from the different sites were harmonized and pooled to increase the power for a joint analysis of the data.

**Table 1:** COHORTs dataset, stratified by site and analysis/hold-out dataset

| SITEID | n<sup>†</sup> | N<sup>‡</sup> | n/N | *Age<sub>min</sub>*<sup>*</sup> | *Age<sub>max</sub>*<sup>*</sup> |
| --- | --- | --- | --- | --- | --- |
| Used in the analysis |  |  |  |  |  |
| Brazil | 9590 | 4158 | 2.3 | 0.7 | 16 |
| Guatemala | 12155 | 1609 | 7.6 | 0 | 18 |
| India | 45584 | 5948 | 7.7 | 0 | 15 |
| Philippines | 31084 | 2451 | 12.7 | 0 | 19 |
| South Africa | 8155 | 2125 | 3.8 | 0.76 | 19 |
| Hold out |  |  |  |  |  |
| Brazil | 2380 | 1048 | 2.3 | 0.7 | 15 |
| Guatemala | 3028 | 392 | 7.7 | 0 | 18 |
| India | 11720 | 1529 | 7.7 | 0 | 15 |
| Philippines | 7612 | 603 | 12.6 | 0 | 19 |
| South Africa | 2163 | 558 | 3.9 | 0.76 | 19 |
| All data |  |  |  |  |  |
| Brazil | 11970 | 5206 | 2.3 | 0.7 | 16 |
| Guatemala | 15183 | 2001 | 7.6 | 0 | 18 |
| India | 57304 | 7477 | 7.7 | 0 | 15 |
| Philippines | 38696 | 3054 | 12.7 | 0 | 19 |
| South Africa | 10318 | 2683 | 3.8 | 0.76 | 19 |

<sup>†</sup> Number of HAZ observations
<sup>‡</sup> Number of children
<sup>\*</sup> Minimal and maximal children ages (in years) for the HAZ observations

**Figure 1:**
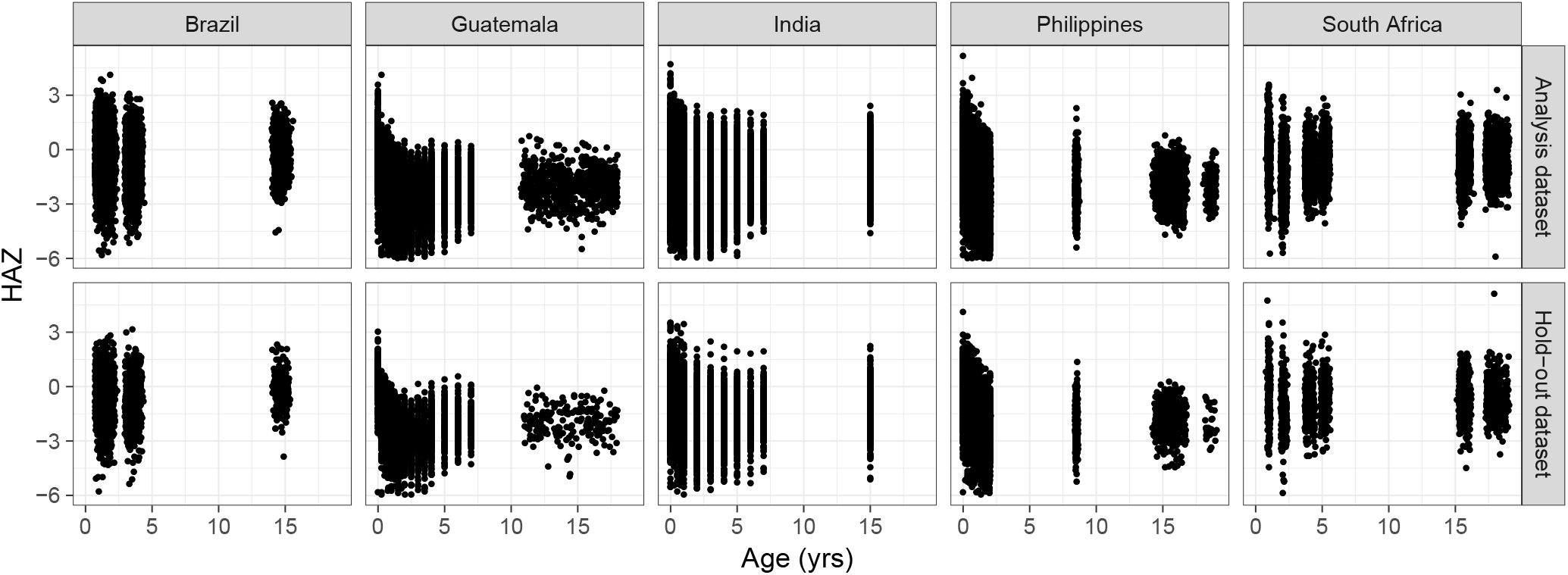
HAZ observations versus age (in years), stratified by site and hold-out versus analysis dataset.

The missing covariate data pattern is visualized in Figure 2 showing that some covariates are missing in some cohorts while others have different levels of missing data for each site. The only covariate that were non-missing (except site) was sex.

**Figure 2:**
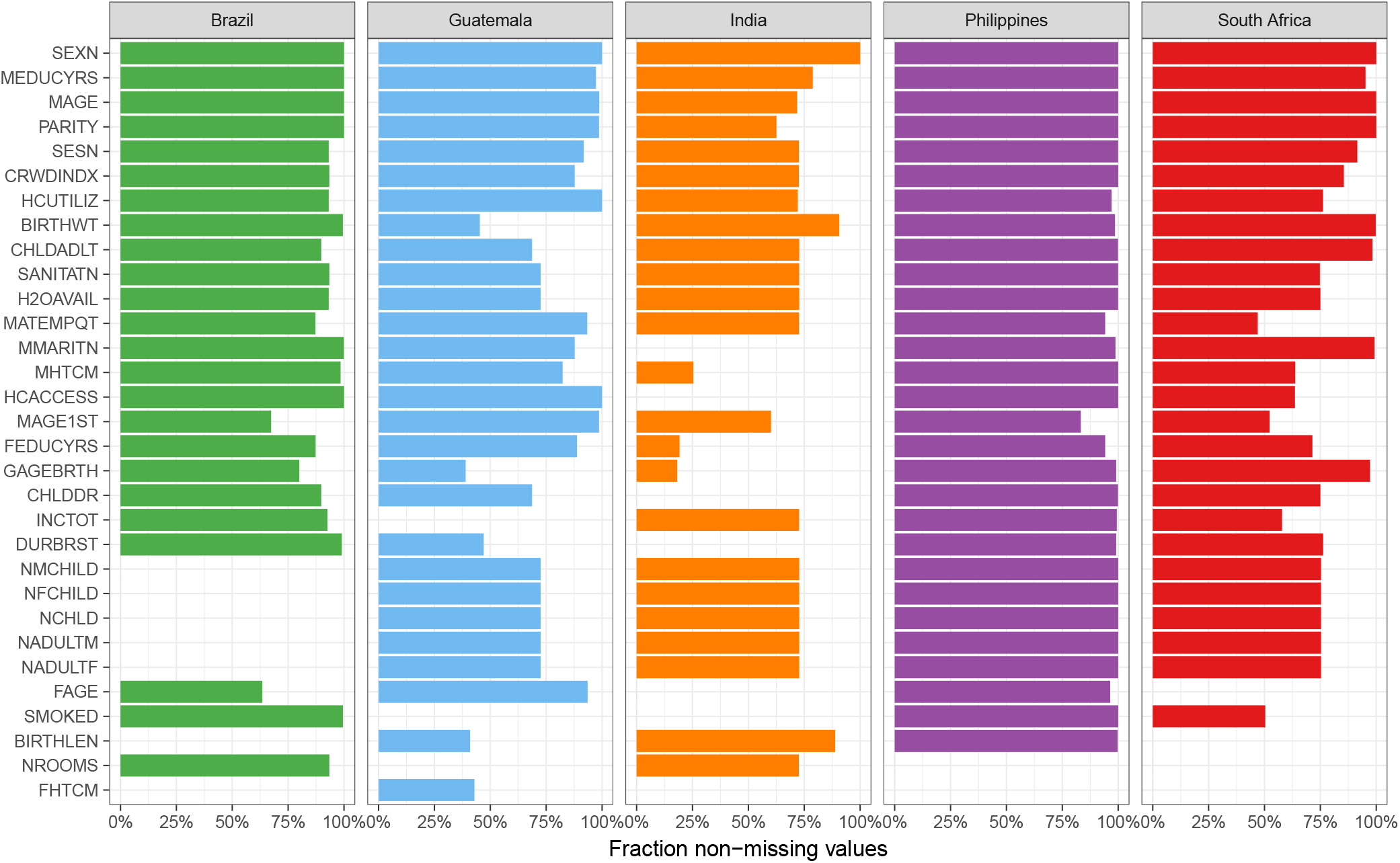
The missing data pattern in the COHORTs study for the covariates used in the model building, stratified by site.

### 3.1 Modeling of the COHORTs HAZ data

A NLMEM was developed (see Section 2) to describe the longitudinal HAZ data. The model building steps are summarized below:

- The data were randomly split into a hold-out dataset of 20% of the children (stratified by site) while the rest of the children where used to build the model.
- A structural model was established, describing the trends of the HAZ data
- Covariate effects where included in the model using the FREM approach, described in Section 2
- Model refinements and predictions into the hold-out data were performed

The covariates that was included in the model were: sex (males/females) - SEXN; maternal years of education - MEDUCYRS; maternal age at birth - MAGE; number of previous births - PARITY; social economic status - SESN; the number of people in house, crowding index - CRWDINDX; use of preventive health care - HCUTILIZ; birth weight - BIRTHWT; all children to adult ratio - CHLDADLT; sanitation - SANITATN; Water availability at water source - H2OAVAIL; maternal empowerment quartile - MATEMPQT; maternal marriage status - MMARITN; maternal height - MHTCM; health care access - HCACCESS; maternal age at first birth - MAGE1ST; paternal years of education - FEDUCYRS; gestational age at birth - GAGEBIRTH; child dependency ratio - CHLDDR; normalized total income in household - INCTOT; duration of breast feeding - DURBRST; maternal number of male children - NMCHILD; maternal number of female children - NFCHILD; number of children in the home - NCHLD; number of male adults in the home - NADULTM; number of female adults in the home - NADULTF; - paternal age at child birth - FAGE; mother smoked during pregnancy - SMOKED; child length at birth - BIRTHLEN; number of rooms in house - NROOMS; fathers height - FHTCM.

The performance of the model was evaluated by VPCs where the model was used to predict the data used to build the model as well as the hold-out dataset. Here, 100 simulated studies where used to compare the model predictions to the observed data.

A NLMEM estimation tool, NONMEM 7.3 (ref) with the importance sampling method (IMPMAP) was used to estimated the FREM model.

## 4 Simulation setup

Three covariates (birth weight (BWT), birth length (BHT), sex (male or female) (SEX)) were used in the simulation study to investigate the missing data performance. The covariates were sampled with replacement (together with the child specific sampling design, on average 8 samples/child from age 0 to 15 years) from the Indian site of the COHORTs dataset. Only children with all three covariates observed (n=6626) were used when sampled from the Indian site. An Indian site version of the COHORTs data were re-estimated (as a FREM model) using the Indian site data only with the three covariates (BWT, BHT, SEX) as predictors. The model parameters are described in appendix XX. The estimated Indian site FREM were re-calculated (see Equation 5) to a FFEM model assuming that all covariates are known. This FFEM was used to simulate new HAZ data.

The setup of the simulation study are described below:

1. Sample children (n=1000 (RICH) or n=100 (SPARSE)) from the COHORTs data with no missing covariate information
2. Simulate data for the sampled children using the simulation FFEM using the observed design for each child
3. Re-estimate using FFEM with the various levels of missing covariates (0%, 10%, 30%, 70% and 90%)
4. Re-estimate using FREM with the various levels of missing covariates (0%, 10%, 30%, 70% and 90%)
5. Transform the estimated FREM to a FFEM for comparison

The procedure were repeated 1000 times to be able to explore bias and precision. All models were estimated using the nonlinear mixed effect models software NONMEM 7.3 using the important sampling (IMPMAP) method.

## 5 Results

### 5.1 The COHORTs model

The structural base model (without covariates) was an inverse Bateman function with a time varying baseline Equation 11. The baseline changed linearly up to an estimated age (*BP*_*i*_) after which it was constant. The slope of the linear change was associated with an additive random effect, meaning that the individual change in baseline could be either positive or negative. The FREM after including covariates described the data well, both in the analysis data set as well as in the hold-out data set (Figure 3).

**Figure 3:**
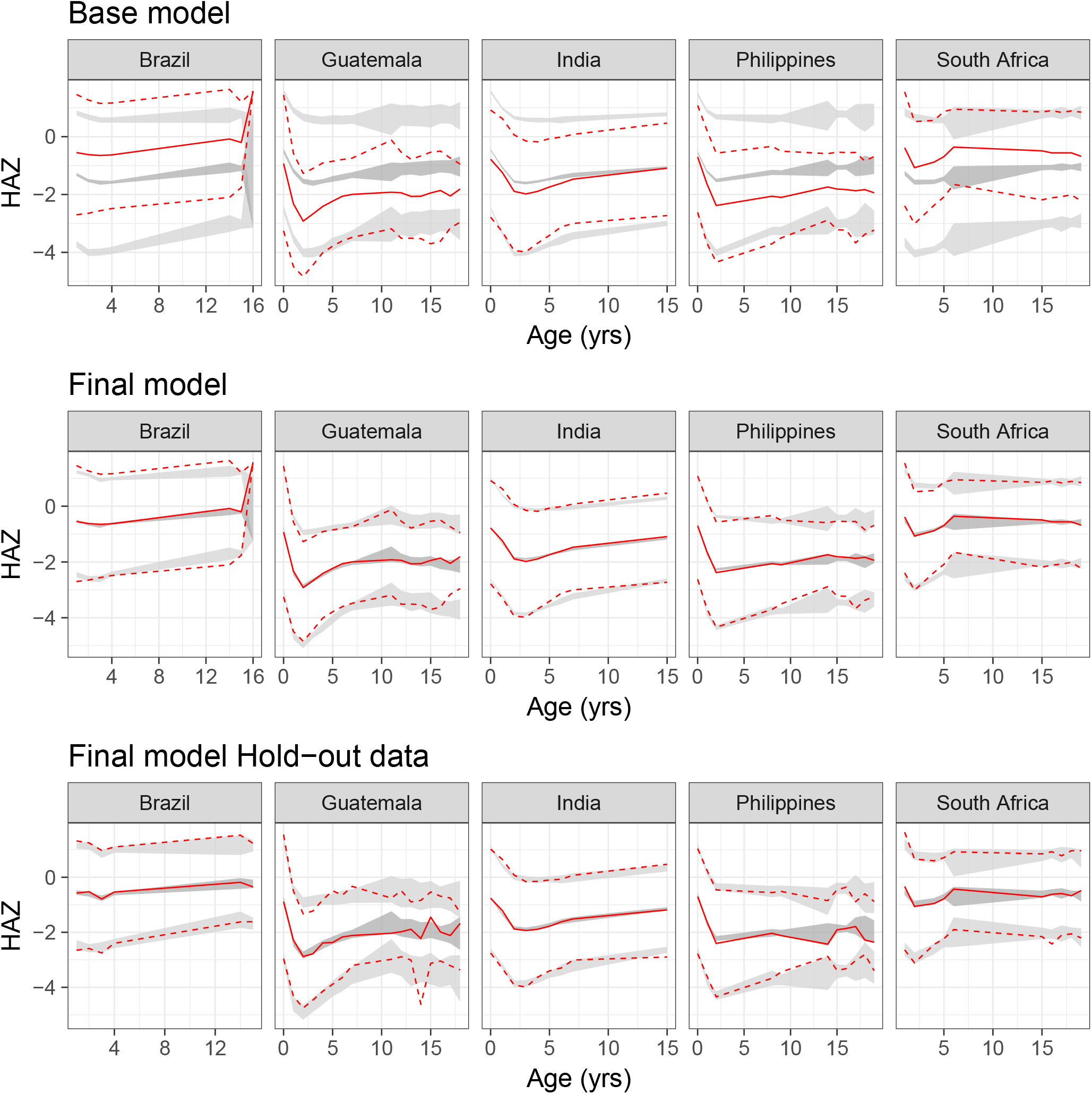
Visual predictive check of HAZ versus age stratified by site. The red lines are the observed 97.5th, 50th and 2.5th percentile of the observed data and the grey shaded areas such that they encompass the central 95% of the corresponding model simulations. The two upper panels are based on the data used to build the model (the base model and the final FREM) while the lower panel are predictions into the hold-out dataset.

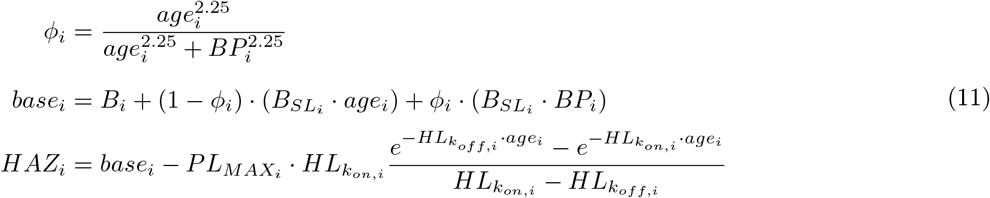

where *age*_*i*_ is the ith child’s observation times (in years since birth). Child specific parameters *B*_*i*_ 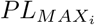 and 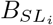 were assumed normally distributed while *BP*_*i*_ 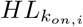 and 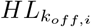 were assumed log-normally distributed, i.e. see Equation 8.

The structural parameter estimates of the final FREM model are presented in Table 2 while all parameters in the final FREM model are presented in the supplement material.

**Table 2:** Parameter estimates of structural part of final FREM model

| Parameter | Typical value | IIV<sup>†,‡</sup> |
| --- | --- | --- |
| $B_i$ | -0.0753 | 1.5181 |
| $B_{SL_i}$ (/year) | -0.0777 | 0.0105 |
| $BP_i$ (year) | 10.4397 | 0.0200 |
| $PL_{MAX_i}$ | 3.3899 | 6.8612 |
| $HL_{kon,i}$ (year) | 1.1522 | 0.3762 |
| $HL_{koff,i}$ (year) | 1.9828 | 0.0735 |

<sup>†</sup> Inter individual variability (on variance scale)

The univariate covariate that explained most variability was the site, followed by sanitation and height of the mother, see Figure 4. This figure is stratified by Nadir (lowest value of each individual prediction) and the predicted HAZ at 2 and 11 years conditioned on each covariate separately. Note that even though the strongest covariate effect (site) explain up to 60% of the explainable variability. The variability that can maximally be explained by the covariates were lower. For HAZ at 2 years, for example, only 22% of the total variability was explained by site.

**Figure 4:**
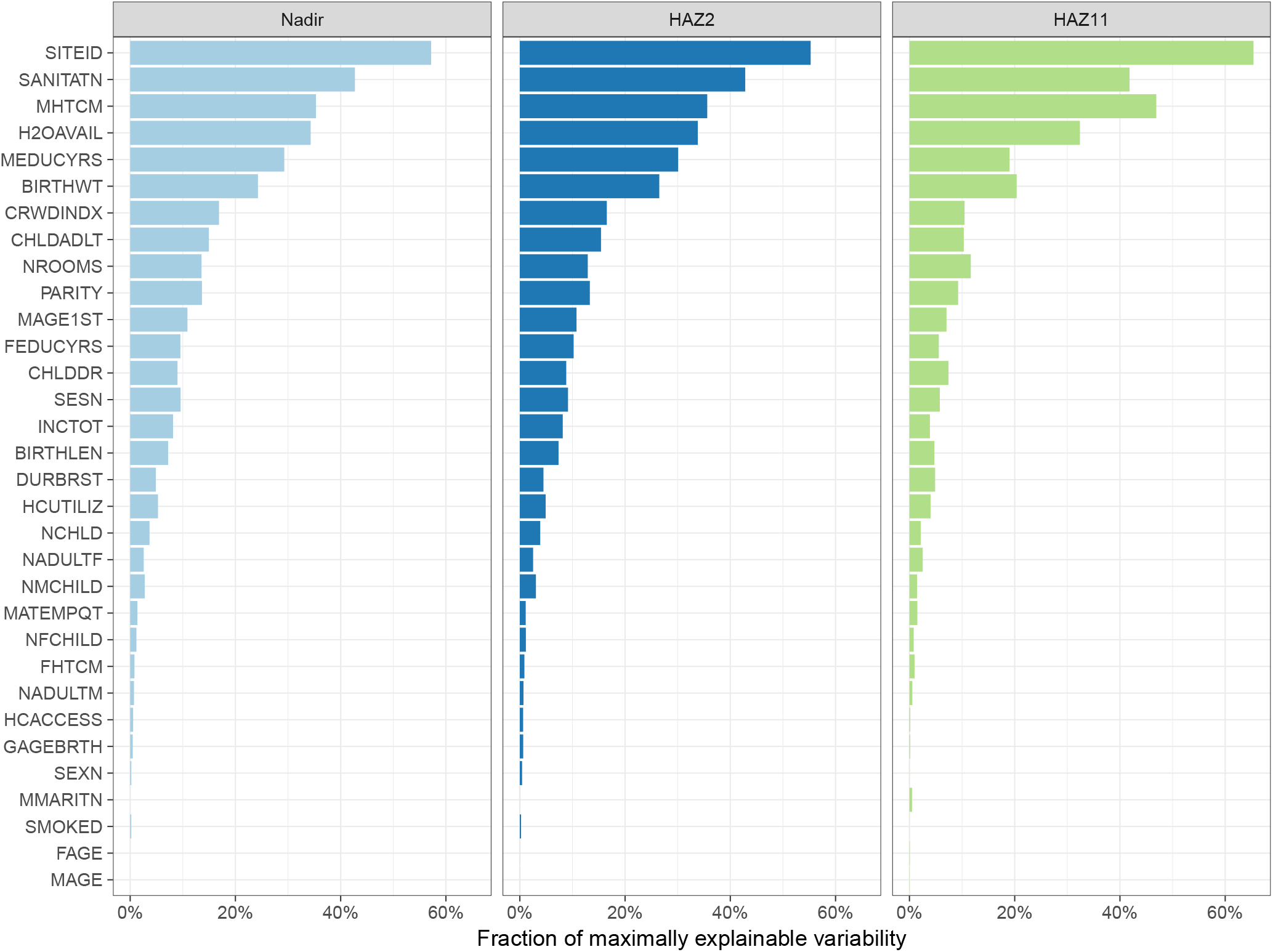
The fraction of the maximally explainable variability in Nadir, HAZ at age 2 years and HAZ at age 11 years that are explained by each of the covariates by them self.

### 5.2 Simulation results

The covariates coefficients, the typical population parameters and the variance estimates are illustrated in Figures 5, 6 and 7 respectively, stratified by level of missingness and NLMEM method. Similar illustrations of the performance with the SPARSE dataset are presented in the supporting information (Section 6).

**Figure 5:**
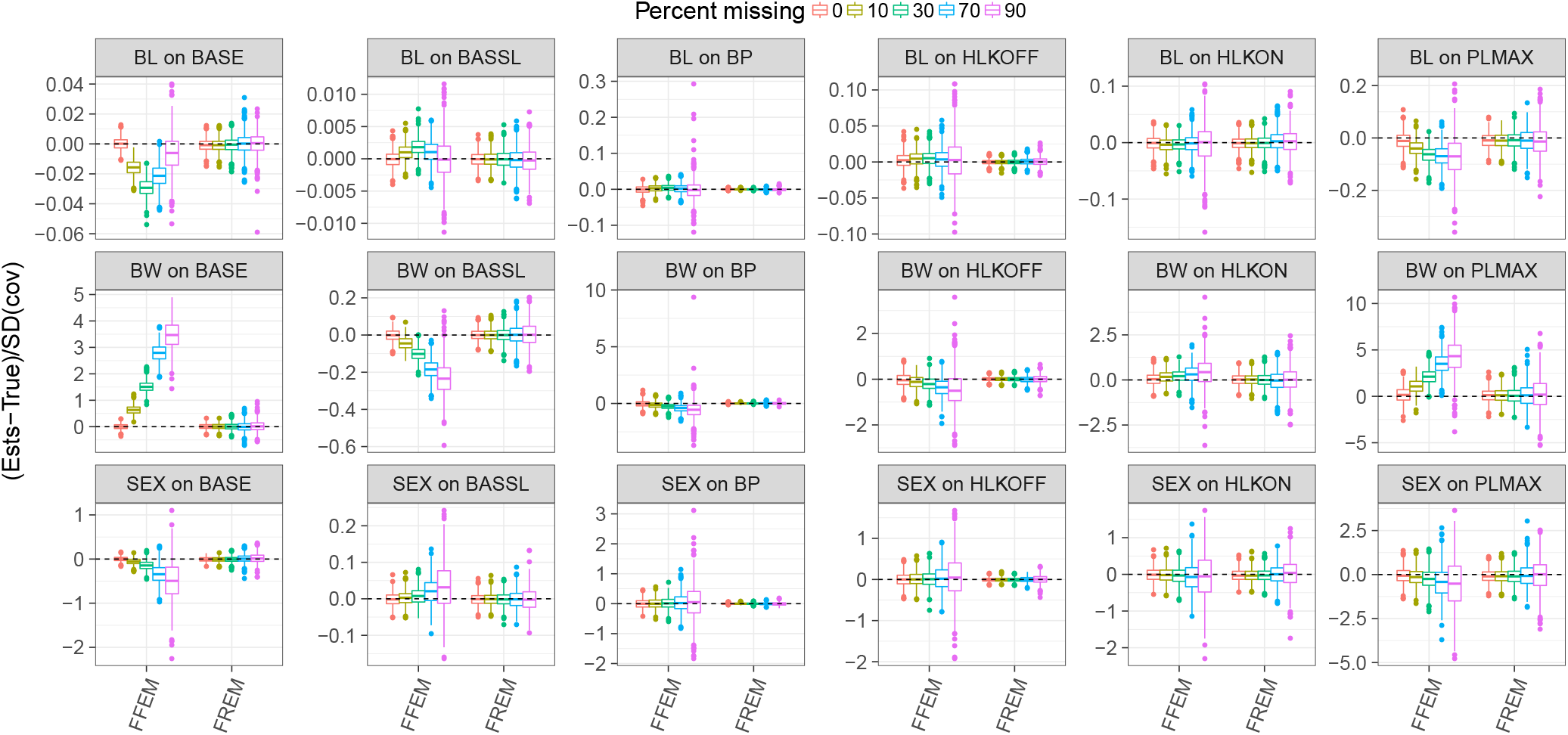
Estimated normalized covariate coefficients with the RICH design, colored by the different levels of missing data. Stratified by modeling technique, FFEM versus FREM.

**Figure 6:**
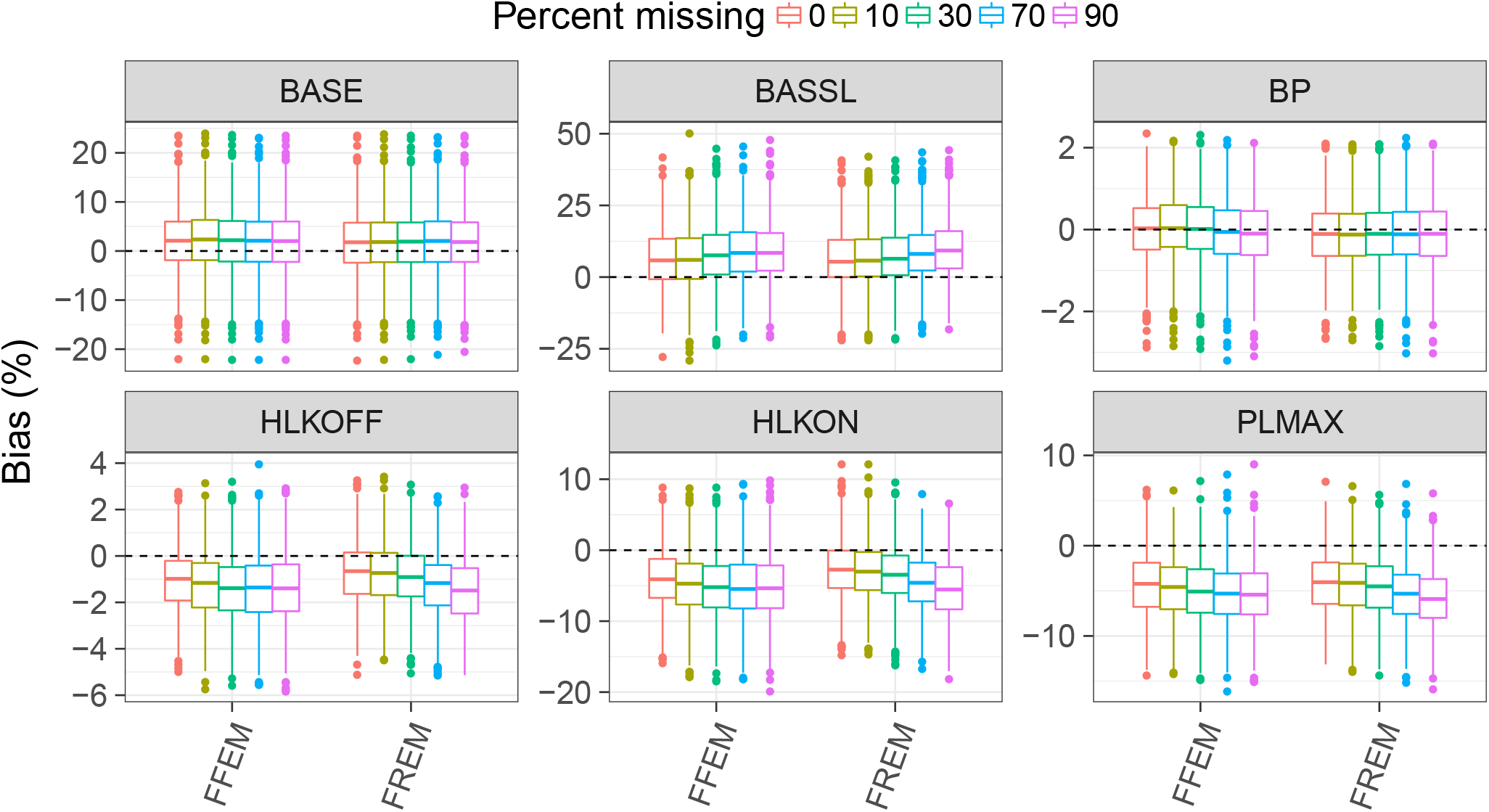
Estimated population typical parameters with the RICH design, colored by the different levels of missing data. Stratified by modeling technique, FFEM versus FREM.

**Figure 7:**
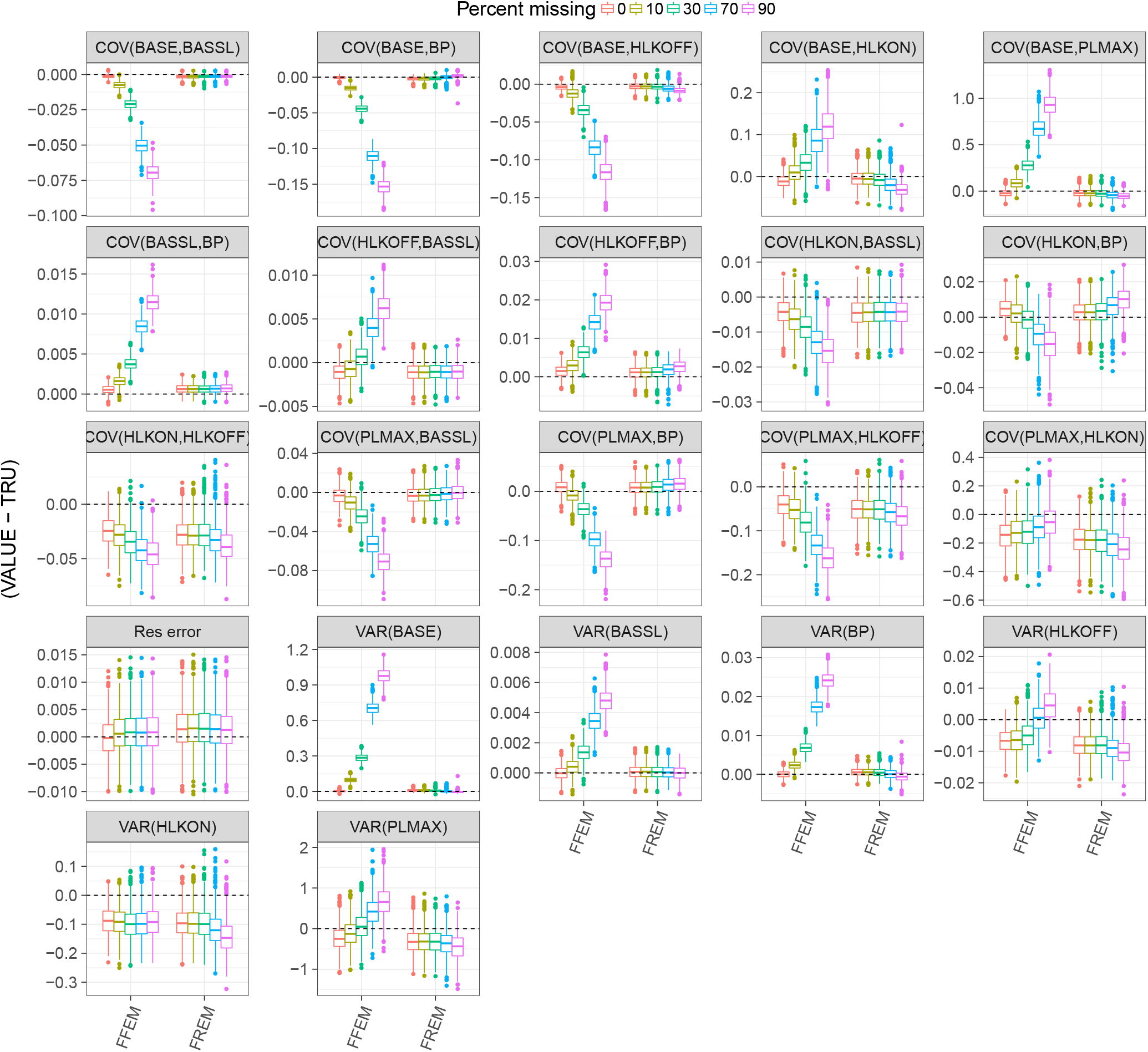
Estimated normalized population variance parameters with the RICH design, colored by the different levels of missing data. Stratified by modeling technique, FFEM versus FREM.

When mean imputation was used with FFEM, the estimated covariate coefficients have, in general, increasing bias with increasing degree of missing covariate information (see Figure 5). This is not the case for FREM which shows more or less unbiased results regardless of the degree of missing covariate data (up to 90%). The precision of the covariate coefficients decrease with the increasing level of missing covariate information, for both FFEM and FREM. This is expected since the missing information directly affect the sample size of the observed covariates. However, the loss in precision with increasing level of missingness is higher for the median imputation with FFEM compared to FREM.

As expected, the structural parameters are almost independent of the degree of missingness (Figure 6). There is a trend towards increasing bias with increasing levels of missingness in both FFEM and FREM but it’s only minor. However, the precision is unaffected by the level of missing covariate information.

The estimated IIV variance-covariance matrix (*D*_*par*_) have increased bias with increasing degree of missing covariate information in most of the variance parameters with FFEM. With FREM, this bias was much smaller and mainly visible in the 90% missing covariate information (see Figure 7). The precision of the variance-covariance matrix is not affected when FREM is used while it decreased with increasing degree of missing covariate information using mean imputation in the FFEM context.

Similar trends as described above, but slightly more pronounced, is seen when sparse data (SPARSE) is used instead. The results for SPARSE are available in the supporting information, Figures 8, 9 and 10, respectively.

**Figure 8:**
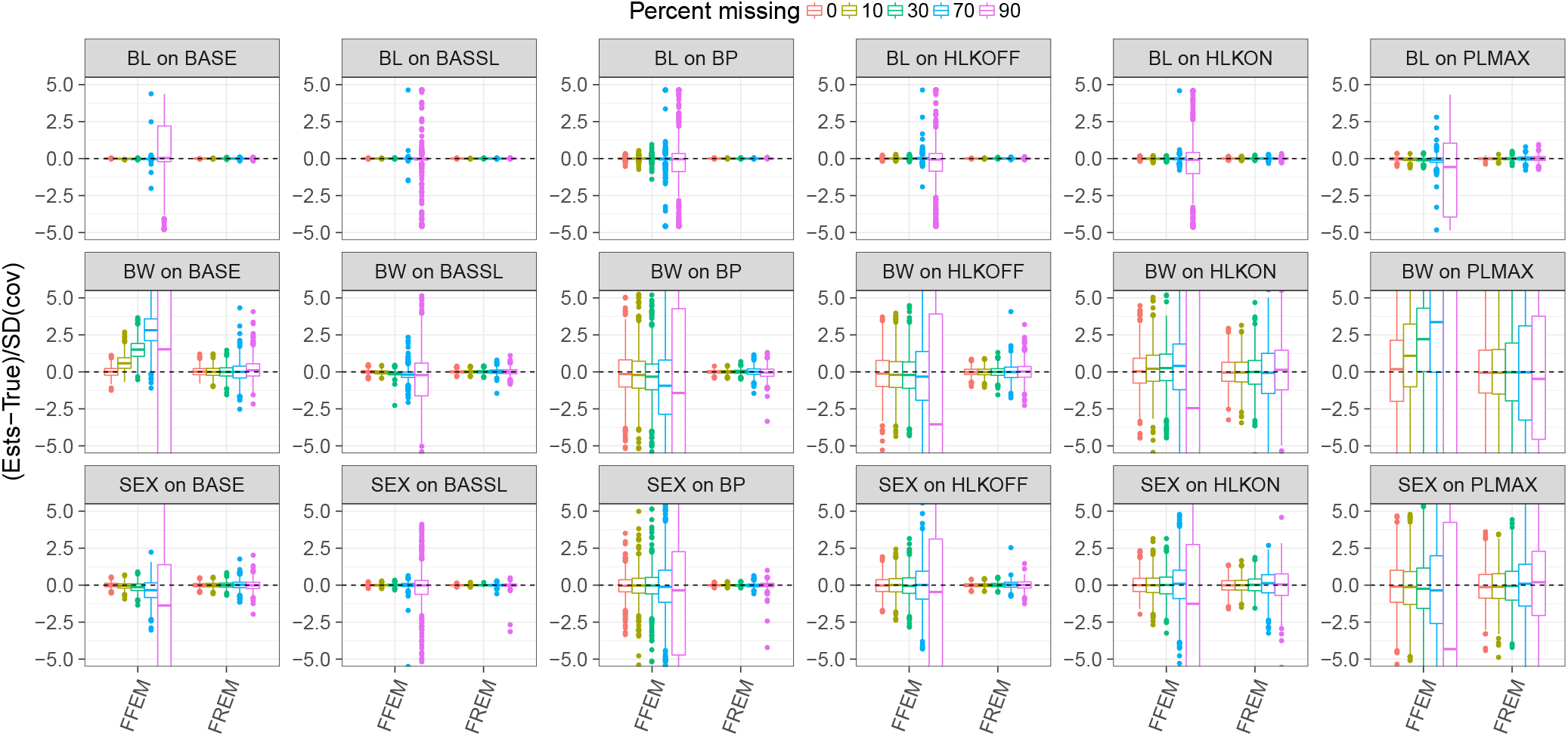
Estimated normalized covariate coefficients with the SPARSE design, colored by the different levels of missing data. Stratified by modeling technique, FFEM versus FREM.

**Figure 9:**
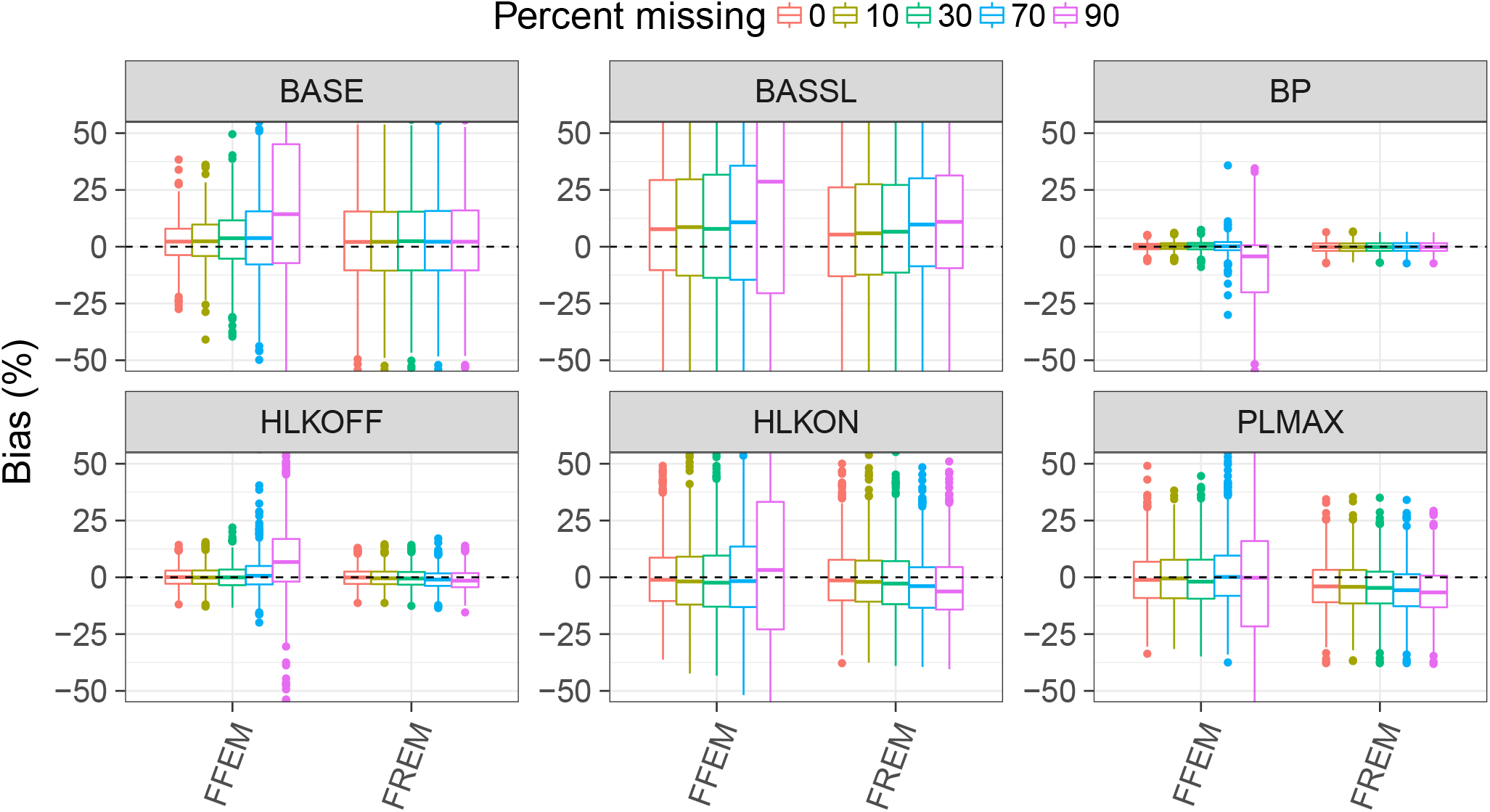
Estimated population typical parameters with the SPARSE design, colored by the different levels of missing data. Stratified by modeling technique, FFEM versus FREM.

**Figure 10:**
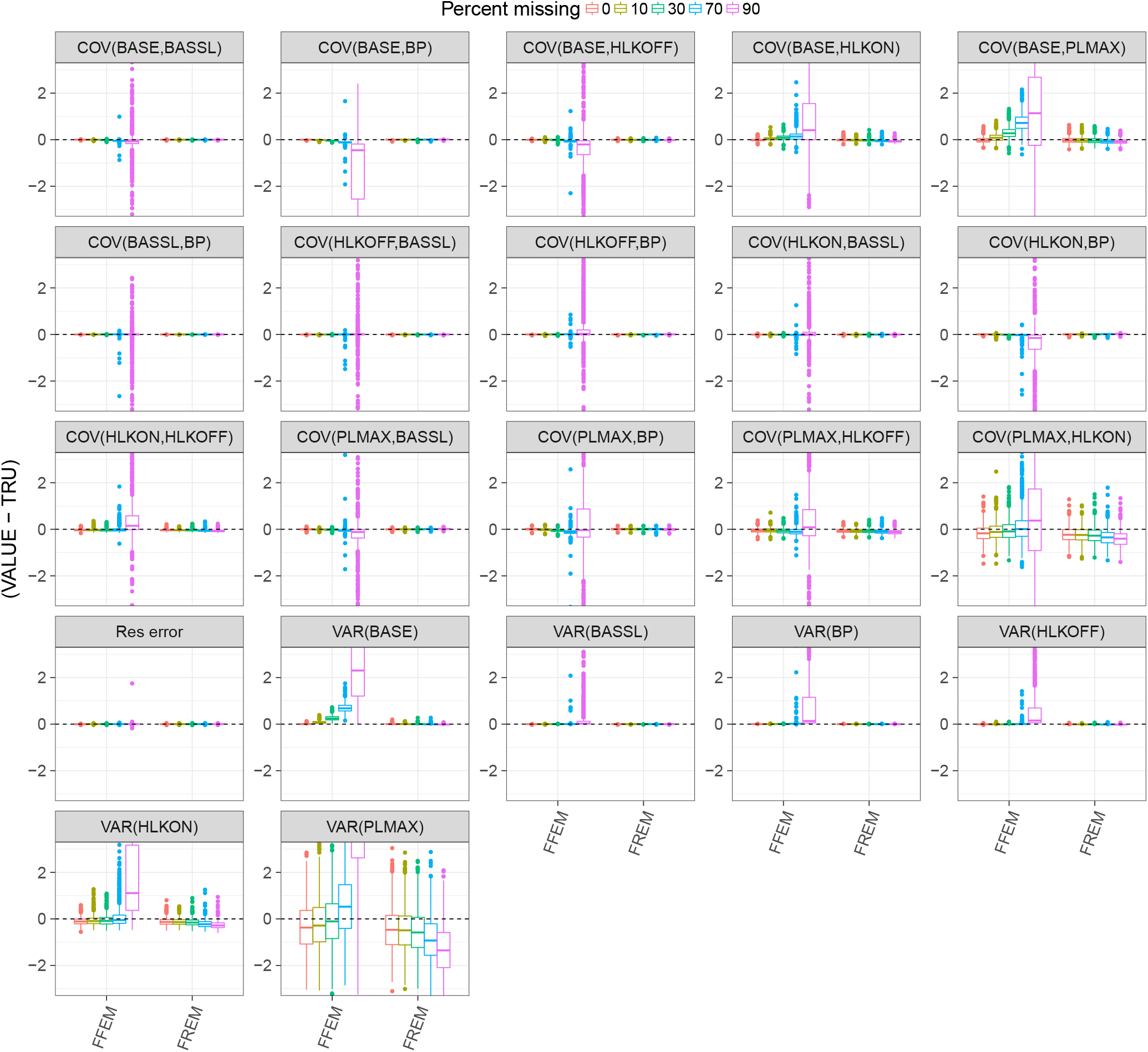
Estimated normalized population variance parameters with the SPARSE design, colored by the different levels of missing data. Stratified by modeling technique, FFEM versus FREM.

## 6 Discussion

A FREM model describing the COHORTs data was developed. The model described the data well and was also able to predict into a hold-out dataset with high predictive performance. This model can therefore be used to design coming studies using either optimal design techniques or simulations assuming that the population in the new study is somewhat similar to the COHORTs population. The analysis identified the most important covariates in the COHORTs study and this information can also be used to design new studies and give further insight into important factors in growth development in low and middle income countries. Nevertheless, the variability in the COHORTs data was large and all covariates together could explain only roughly 22% of the variability in HAZ at 2 years, HAZ at nadir and HAZ at 11 years. This indicates that it is important to further investigate additional covariates that can help identifying the most important factors that influence physical growth. The low explainable variability also opens up for performing studies where designs are optimized for each individual, similar to ideas in precision medicine. The FREM model and in general NLMEM are very suitable for using the model as prior information and optimize individual designs and potentially interventions. Moreover, the FREM model can be transformed to predict designs using any combination of covariates that are included in the model and therefore suitable for new designs where only part of the FREM included covariates are available for inclusion. Site was the covariate that explained most variability in the FREM model, however site is potentially a changing covariates since e.g. Brazil 50 years ago might be very different from Brazil today and similarly; different regions in Brazil might have very different outcomes when it comes to physical growth. Hence, using the model to extrapolate to other sites might need the adjustment on how a site (or country) has developed since the COHORTs study.

FREM appears to be suitable as an approach for handling missing data. The main reason for this is that FREM compensates for the missing information via the correlation between the full block of IIV with both the non-missing covariates and the parameters. Also compared to other methods, e.g. multiple imputation (5), the handling of missing data is implicit in FREM, hence no additional overhead is added when handling missing data. Even though the distribution of the missing data were misspecified (e.g. SEX is not normally distributed), the FREM handling of missing data gives reasonable precise and unbiased estimates even with 90% missing covariate information. This is not the case with mean imputation using a FFEM.

In the current analysis the ability of the FREM approach to handle missing data was compared to simple mean imputation. Alternative FFEM based methods for NLMEM analyses, such as multiple imputation and likelihood based methods, are due to their complex implementation in NLMEM not particularly common (although there are a few examples REF) and was therefore not included in the comparison. It is expected, though, that they would have performed better than mean imputation and probably on par with the FREM approach.

To conclude; FREM improves the estimation properties when handling missing data compared to median single imputation with FFEM. In both the RICH and SPARE dataset, FREM estimated coefficients were unbiased, even when 90% missing data. This suggests that FREM is an appropriate method to use for handling situations with missing covariate data such as in global health studies, especially since missing covariates are automatically handled within FREM.

## Acknowledgments

This study was supported by the Bill & Melinda Gates Foundation. The article contents are the sole responsibility of the authors and may not necessarily represent the official views of the Bill & Melinda Gates Foundation or other agencies that may have supported the primary data studies used in the present study. The authors wish to recognize the members of the HBGDki community, including the principal investigators and their study team members for their generous contribution of the data that made this report possible and the members of the HBGDki team who directly or indirectly contributed to the study.

## Author contributions

This is an author contribution text. This is an author contribution text. This is an author contribution text. This is an author contribution text. This is an author contribution text.

## Supporting information

The following supporting information is available as part of the online article:

The covariates coefficients, the typical population parameters and the variance estimates for the SPARSE data scenario are illustrated in Figures 8, 9 and 10 respectively, stratified by level of missingness and NLMEM method.

## References

1. Gastonguay. Marc R. Full Covariate Models as an Alternative to Methods Relying on Statistical Significance for Inferences about Covariate Effects: A Review of Methodology and 42 Case Studies. PAGE 20 (2011) Abstr 2229 [www.page-meeting.org/?abstract=2229].

2. Rubin Donald B. Inference and missing data. Biometrika. 1976;63(3):581–592, DOI 10.1093/biomet/63.3.581, available at /oup/backfile/content_public/journal/biomet/63/3/10.1093/biomet/63.3.581/2/63-3-581.pdf.

3. Hu C. Zhang J. and Zhou. Confirmatory analysis for phase III population pharmacokinetics. Pharm Stat. 2011;10(1):14–26.

4. Hu C. and Zhou H. An improved approach for confirmatory phase III population pharmacokinetic analysis. J Clin Pharmacol. 2008;48(7):812–822.

5. Rubin Donald B. Multiple imputation for nonresponse in surveys; 1987.

6. Novakovic A. M. and Krekels. Application of Item Response Theory to Modeling of Expanded Disability Status Scale in Multiple Sclerosis. AAPS J. 2017;19(1):172–179.

7. Brekkan A. and Lopez-Lazaro. A Population Pharmacokinetic-Pharmacodynamic Model of Pegfilgrastim. AAPS J. 2018;20(5):91.

8. Abrantes J. and Solms. Integrated modelling of factor VIII activity kinetics, occurrence of bleeds and individual characteristics in haemophilia A patients using a full random effects modelling (FREM) approach (www.page-meeting.org/?abstract=8646). 2018.

9. M. O. Karlsson. A full model approach based on the covariance matrix of parameters and covariates (www.page-meeting.org/?abstract=2455); 2012

10. Richter L. M. and Victora. Cohort profile: the consortium of health-orientated research in transitioning societies. Int J Epidemiol. 2012;41(3):621–626.

